# Detection of somatic structural variants from short-read next-generation sequencing data

**DOI:** 10.1101/840751

**Authors:** Tingting Gong, Vanessa M Hayes, Eva KF Chan

## Abstract

Somatic structural variants (SVs) play a significant role in cancer development and evolution, but are notoriously more difficult to detect than small variants from short-read next-generation sequencing (NGS) data. This is due to a combination of challenges attributed to the purity of tumour samples, tumour heterogeneity, limitations of short-read information from NGS, and sequence alignment ambiguities. In spite of active development of SV detection tools (callers) over the past few years, each method has inherent advantages and limitations. In this review, we highlight some of the important factors affecting somatic SV detection and compared the performance of eight commonly used SV callers. In particular, we focus on the extent of change in sensitivity and precision for detecting different SV types and size ranges from samples with differing variant allele frequencies and sequencing depths of coverage. We highlight the reasons for why some SV callers perform well in some settings but not others, allowing our evaluation findings to be extended beyond the eight SV callers examined in this paper. As the importance of large structural variants become increasingly recognised in cancer genomics, this paper provides a timely review on some of the most impactful factors influencing somatic SV detection and guidance on selecting an appropriate SV caller.

## Introduction

Cancer is a disease of the genome that develops through the accumulation of somatic mutations (variants), ranging from single nucleotide variants (SNVs), insertions/deletions (indels) of a few nucleotides, to large structural variants (SVs) [1]. SVs are typically defined as genomic rearrangements involving at least fifty nucleotide bases (50 bp), and broadly include deletions, insertions, duplications, inversions and translocations [2]. SV formation can leave complex genomic patterns, some of which are associated with specific cancer types while others are more broadly implicated. For example, chromothripsis, involving hundreds of clustered rearrangements arising from the shattering and inaccurate reassembly of a single chromosome, are found in 3% of all cancer types [3]. In contrast, chromoplexy, involving complex coordinated chains of rearrangements are almost exclusively found in prostate cancer [4]. Thus, the extent and types of somatic SVs can help characterize tumour types and provide insights into the mechanisms of oncogenesis. Additionally, SVs that underlie oncogene activation and tumour suppressor loss, can potentially be targeted for therapy or used as prognostic markers [1].

Due to continued reductions in cost and increase in throughput, next-generation sequencing (NGS) has become the preferred approach for cancer genomics [1,5]. NGS, currently dominated by Illumina pair-end short-read sequencing, is a technology that allows the entire human genome to be read. Conceptually, the approach is simple: DNA from multiple cells is extracted and fragmented to a desired library size (typically 200-500 bp), then the ends of each fragment are tagged (so they can be tracked and paired) and sequenced inwards to up to ∼150 bp [6]. Following sequencing, the billions of reads are informatically paired and aligned to a known reference genome for variant detection. SVs are inferred from abnormal alignment patterns suggestive of genomic rearrangement breakpoints. The underlying bioinformatic analyses is not straightforward for several reasons. Detection of SVs that are kilobases to megabases in length is difficult from short sequence reads and small insert library sizes (distance between pairs of reads) as they cannot be captured by any single sequence [7]. While sequencing of more reads (higher depth of coverage) can sometimes compensate for this, it provides limited advantage at genomic regions with low sequencing complexity (e.g. repetitive sequences) or regions of high sequence similarity (e.g. segmental duplicated regions). These regions can lead to ambiguous read alignments, which are a significant source of false positive variant detection.

Compared to germline SVs, the detection of somatic SVs in cancer genome (identification of SVs present in the cancer sample but absent in the patient matched normal sample) is further complicated by tumour purity (fraction of cancerous to normal cells) and tumour heterogeneity (presence of clonal and subclonal tumour cell populations). Compounding onto this is that, the extents of both of these confounders are typically unknown at the time of tissue sampling. Again, while increasing sequencing coverage can assist in capturing low abundance tumour SVs, in many cases, it is unclear whether the associated increase in cost can outweigh any information gained [7].

These challenges have resulted in the development and refinement of multiple SV detection methods and SV calling software (SV callers) in the last decade, each with their advantages and disadvantages. While studies have explored some of these effects on SV detection, many have focused on germline SVs [8,9]. The most comprehensive study with focus on somatic SV detection compared the performance of 13 SV callers on three sets of simulated data [10]. They observed that some variables generated similar error profiles across SV callers but did not delve into the extent of these correlations.

In this paper, we provide a comprehensive review on common factors affecting the identification of somatic SVs and on the performance of eight commonly used SV callers. Specifically, we evaluate and quantify each SV caller’s ability to detect different SV types and size ranges, the individual and interaction effects of SV abundance and sequencing coverages, their precision in predicting genomic breakpoints, and the impact of sequence similarity (genomic segmental duplications) on somatic SV detection.

## Methods

To objectively assess the impact of each parameter (SV type, SV size, variant allele fraction, sequencing coverage, sequence similarity) on different SV callers, we used a simulation framework detailed in **Supplementary Methods**. In brief, for each evaluation setting (reviewed in the following sections), three replicate pairs of normal and tumour genomes were simulated to contain germline-only and germline plus cancer SVs, respectively, based on the human reference genome sequence GRCh38. Each simulated SV contains 1200 SVs including 200 of each of six SV types as described in the “Structural variant types and definitions” section. Paired-end short-read sequences were then sampled from the augmented genomes using SVEngine [11] to the desired coverage. SV detection was performed following standard read-alignment against GRCh38 using BWA-MEM v0.7.17-r1194 [12]. Sensitivity and precision of SV callsets were evaluated based on two true positive criteria: (1) the SV type reported for a candidate SV must match the simulated SV, and (2) the genomic position of the reported breakpoints must be within a pre-defined distance from the simulated SV. Unless otherwise stated, evaluation results presented in this study are based on the default breakpoint-resolution threshold of 200 bp as used in similar studies [8,9].

## Results

### Structural variant detection methods and callers

There are four main methods for the detection of SVs from short-read NGS data **(Table 1)**: read-pair, read-depth, split-read, local-assembly [13]. Each of these methods are reliant on pre-alignment of sequencing reads to a reference genome.

**Table 1.**
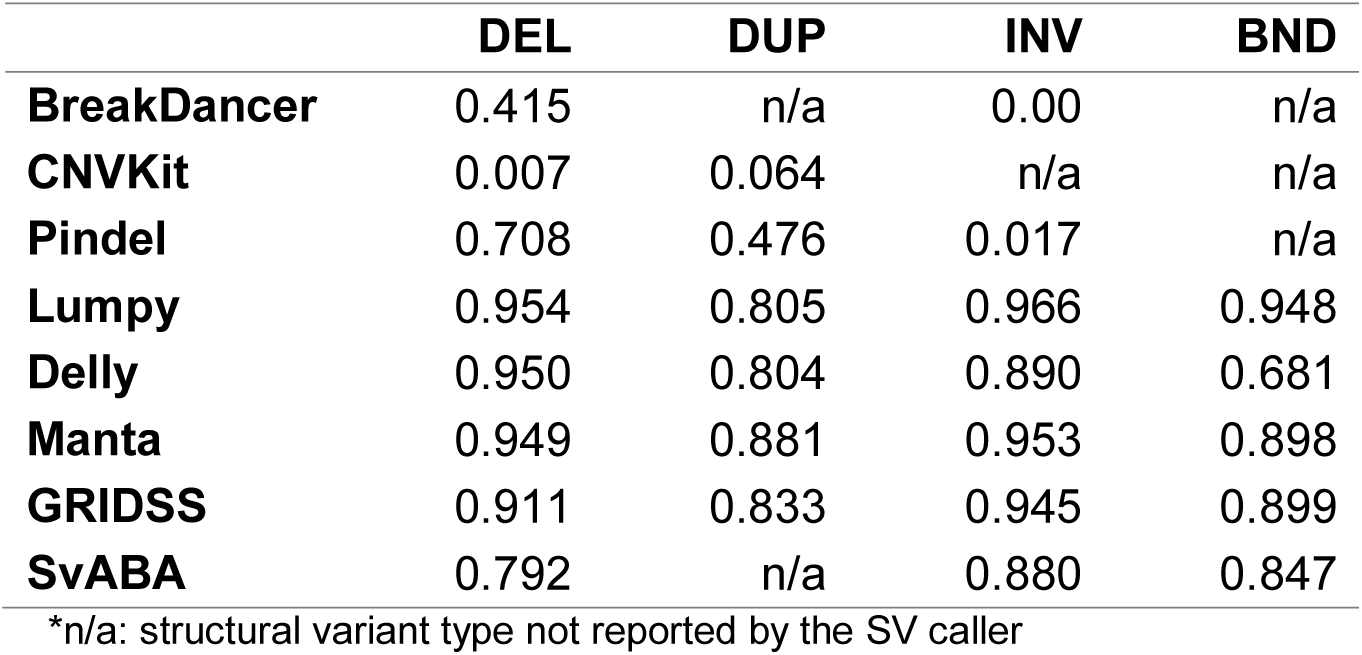
SV detection methods and example SV callers.

The read-pair method searches discordant alignment signatures of paired-end sequencing reads. SVs are identified from read-pairs whose mapped interval is different from the sequencing library insert size or mapped in abnormal orientation. Compared to other methods (such as split-read), read-pair is less sensitive to small insertions and deletions, especially SVs smaller than the library insert size [2]. BreakDancer is an example SV caller that uses the read-pair method [14].

The read-depth method seeks changes in the amount of sequencing coverage at a given genomic interval (segment window) relative to neighbouring or genome-wide coverage. This method is particularly suited for the detection of copy number changes but the resulting breakpoint resolution is poor and heavily dependent on overall sequencing depth and the size of the segment window. In addition, this method can only detect duplications (copy number gains) and deletions (copy number losses). CNVkit is an example SV caller that uses the read-depth method [15].

The split-read method assesses continuity and completeness of NGS read alignments against a reference genome. Discontinuous and incomplete read alignments are indicative of SV events, from which breakpoints can be inferred to single nucleotide resolution. However, while an incomplete alignment may signal the presence of a rearrangement breakpoint, the unaligned portion provides little information on the adjacent sequence nor the type and size of the SV. This is particularly problematic for large insertions; a distinct disadvantage from not using paired-end information. In addition, genomic regions with high sequence similarity can confound the split-read method, and high sequencing depth is often needed to obtain sufficient split-reads overlapping the breakpoint to achieve a confident SV call. Pindel is an example of (and the first) SV caller to use the split-read method; it was one of the tools used to generate the callset in the 1000 Genome Project [16,17].

The local-assembly method attempts to reconstruct rearranged genomic sequences by assembling sequencing reads associated with SV breakpoints as determined from the initial read-alignment. Therefore, local-assembly is usually implemented along with read-pair and split-read methods. Significant advantages of this approach are its ability to detect SVs at genomic regions with higher levels of sequence identity (non-uniqueness) and its ability to reconstruct novel inserted sequences and small highly rearranged sequences. However, a key disadvantage is the requirement for sufficient variant reads for reliable consensus contig assembly; a potential problem for cancer genomes where variants may only be present at low levels. GRIDSS [18] and SvABA [19] are example SV callers utilising genome-wide local assembly.

Aiming to overcome inherent limitations and to take advantage of the different approaches, a number of SV callers have incorporated multiple methods. For example, Delly [20] and Lumpy [21] call SVs using both discordant paired-end and split-read alignments. Delly predicts SVs using discordant paired-end reads then use split-reads to refine SV breakpoints. In contrast, Lumpy integrates multiple alignment signatures into a single SV discovery process. In addition to paired-end and split-read alignment signatures, Manta [22], GRIDSS and SvABA further incorporate local-assembly to improve SV detection. Manta assembles reads in candidate SV regions identified from discordant paired-end and split-read alignments (i.e. targeted assembly), to validate and refine breakpoints, while GRIDSS and SvABA assemble all aberrantly aligned reads to identify breakpoints. SvABA applies assembly in local 25-kbp assembly windows, called windowed local assembly, while GRIDSS performs assembly of all reads aligned improperly and terms it genome-wide break-end assembly.

There are currently hundreds of SV callers available for SV detection from NGS data. They are mainly designed for the detection of germline SVs, which are variants relative to the genome reference. In contrast, the identification of somatic SVs must exclude those observed in both the tumour and matched-normal samples as they represent either germline SVs (hence not relevant to the cancer genome) or are reference artefacts [23]. Most somatic SV callers perform two-sample (matched tumour-normal pair) variant calling, and require further manual filtering (typically by the user) for SV calls present only in the tumour sample (e.g. BreakDancer, Lumpy, GRIDSS). For some SV callers, such as Manta, Delly and SvABA, the filtering step is automated.

In general, SV callers leveraging multiple detection methods have the best balance between sensitivity and precision for the detection of germline SVs, though there are notable differences in their performance for different SV types and sizes [8,9]. For somatic SVs, the recent ICGC-TCGA DREAM Somatic Mutation Calling Challenge, which evaluated the performance of 13 SV callers, found the overall sensitivity and precision of somatic SV calling to be highly influenced by lower allelic fractions of subclonal variants, tumour sequencing depth and read-alignment quality at SV breakpoints [10]. However, this study did not include other important factors such as SV types and sizes.

#### SV callers based on multiple methods are more reproducible

In this section, we compare the overall performance of the eight SV callers representing the common SV detecting methods (**Table 1**). As the SV callers are reliant on different detection methods, their predicted SVs are rarely completely concordant and the total number of somatic SV calls can be widely different (**Figure 1**). As previously noted, Pindel is very sensitive to split-read signatures, thus the majority (97.7%) of detected variants are < 50bp, which are typically considered as small indels rather than large SVs. For the purpose of this study, variants < 50 bp detected by Pindel were excluded. Overall, Pindel, CNVKit and BreakDancer, which are based on a single SV detection method, detect many unique calls, not identified by other callers. In contrast, SV callers based on at least two SV detection methods, are more concordant. Comparing their overall performance (**Figure 2 green bars**), we confirmed SV callers based on more than one SV detection methods have higher sensitivity and lower false discovery rate, similar to observations for germline SV detection [8,9]. Additionally, these SV callers (Manta, Lumpy, GRIDSS, SvAVA and Delly) were observed to have higher precision (> 90%) than sensitivity, ranging from 63% (SVABA) to 89% (Manta).

**Figure 1.**
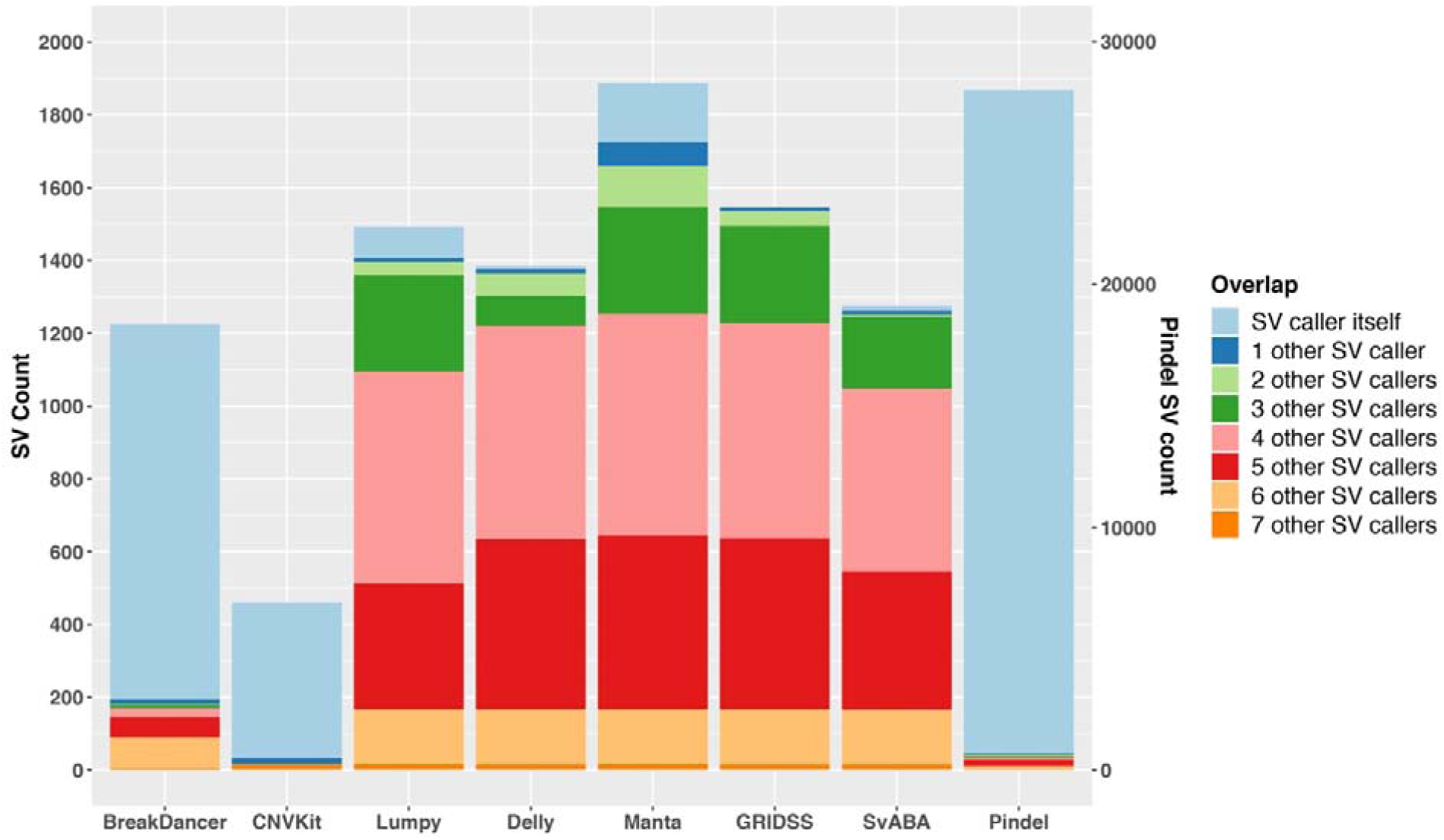
Extent of structural variants identified by different callers and their overlaps. Results are based on simulation data with tumour and matched normal coverage of 60x and variant allele fraction of 100%. The count of Pindel is labelled on the right vertical axis.

**Figure 2.**
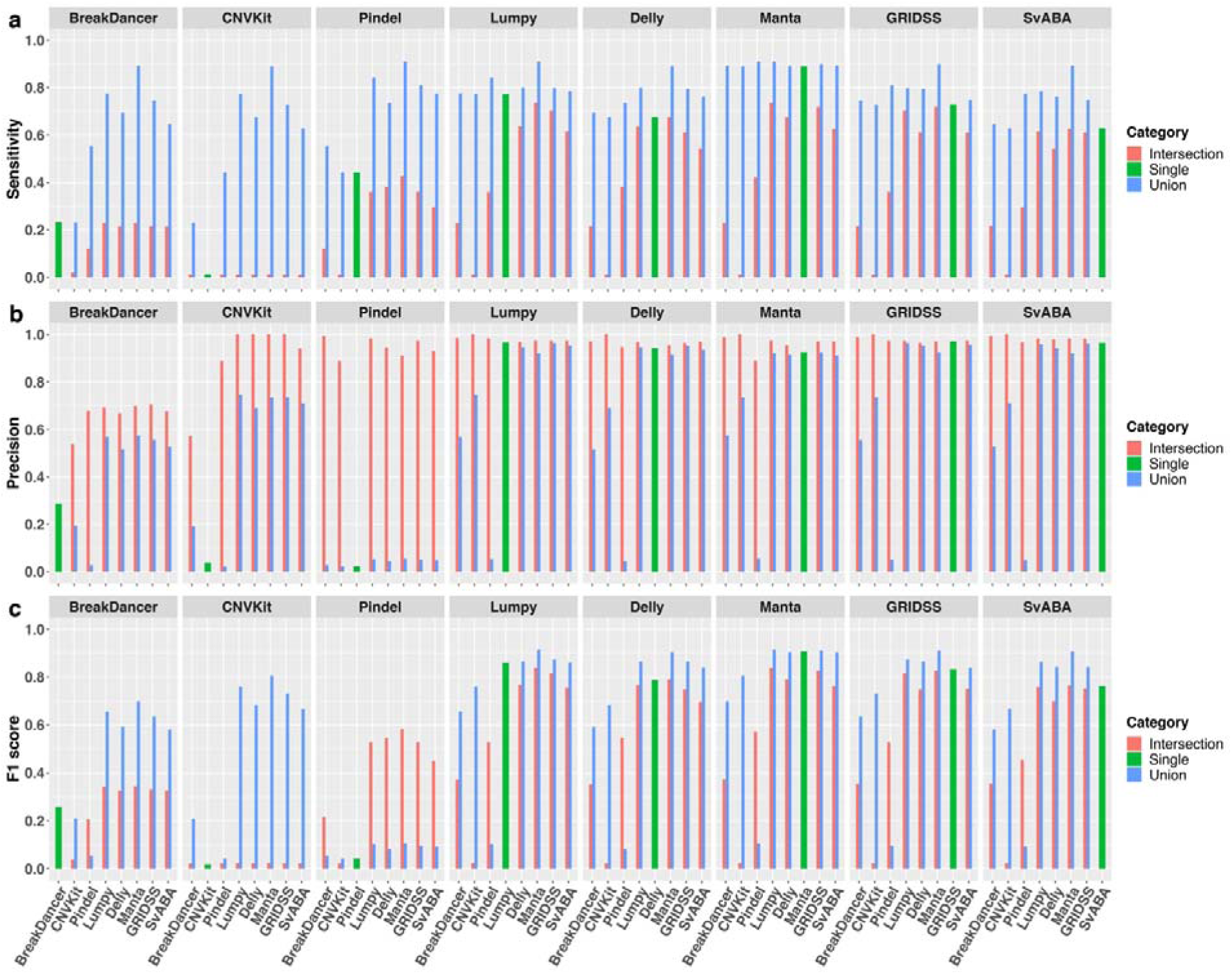
Overall sensitivity, precision and F1 score of individual SV callers and their pairwise union and intersection callsets. Results are based on simulation data with tumour and matched-normal coverage of 60x, variant allele fraction of 100% and breakpoints within 200 bp of simulated SVs.

#### Inclusion of additional callers to an already high-performing SV caller provides little gain to sensitivity and precision

Due to inherent limitations of each SV caller, it is common practice to use at least two SV callers [13] in order to maximise detection sensitivity and precision. In general, a union callset from a pair of SV callers will be more sensitive than either of the single SV callers alone but at the cost of reduced precision, while an intersection callset will improve precision at the cost of reduced sensitivity. To evaluate the extent of gain when combing SV callers, we compared the performance of all pairs of SV callers for both union and intersection callsets. As expected, union callsets (**Figure 2 blue**) greatly improve sensitivity (> 14%), compared to individual SV callers, with the exception of Manta, which has at most only 2% increase when combined with Lumpy or Pindel. However, the impact of union callsets on precision is more variable. When coupled with an “imprecise” SV caller (precision < 30% as a standalone SV caller), such as BreakDancer, Pindel or CNVKit, precision of the union callsets can be worsen by up to 40%. In contrast, the union callsets of two similarly “precise” SV callers (precision > 90%) typically yield little gain on precision (< 5%) compared to using just a single caller. As expected, we observed dramatic improvement in precision when the callset of an imprecise SV caller is intersected with a precise caller, while less improvement (< 10%) is observed when both callers in the pair are already “precise”. In contrast, changes in sensitivity of intersection callsets are more variable. For example, the intersection callsets between Manta and Lumpy or BreakDancer improved precision by 5% and 6% respectively, compared to using Manta alone, but reduce sensitivity by 15% and 66% respectively. Overall, Manta achieved the highest F1 score among individual SV callers and, at most, only about 1% increase in F1 score was gained through its union callsets with Lumpy or GRIDSS (**Figure 2c**).

#### Better performing SV callers do not require longer run time

In addition to overall performance in somatic SV detection accuracy, the amount of computational resources required to run any bioinformatics software can pose logistical limitations. As such, we also compared the CPU time for each of the eight SV callers (**Figure 3**). Pindel was found to have the longest run time, as previously observed [8,9]. The pre-processing step of Lumpy using SAMTOOLS (detail in **Supplementary Methods**) was found to require a large amount of CPU time. In addition, SV callers incorporating local-assembly typically require more CPU time, though Manta appears to be an exception.

**Figure 3.**
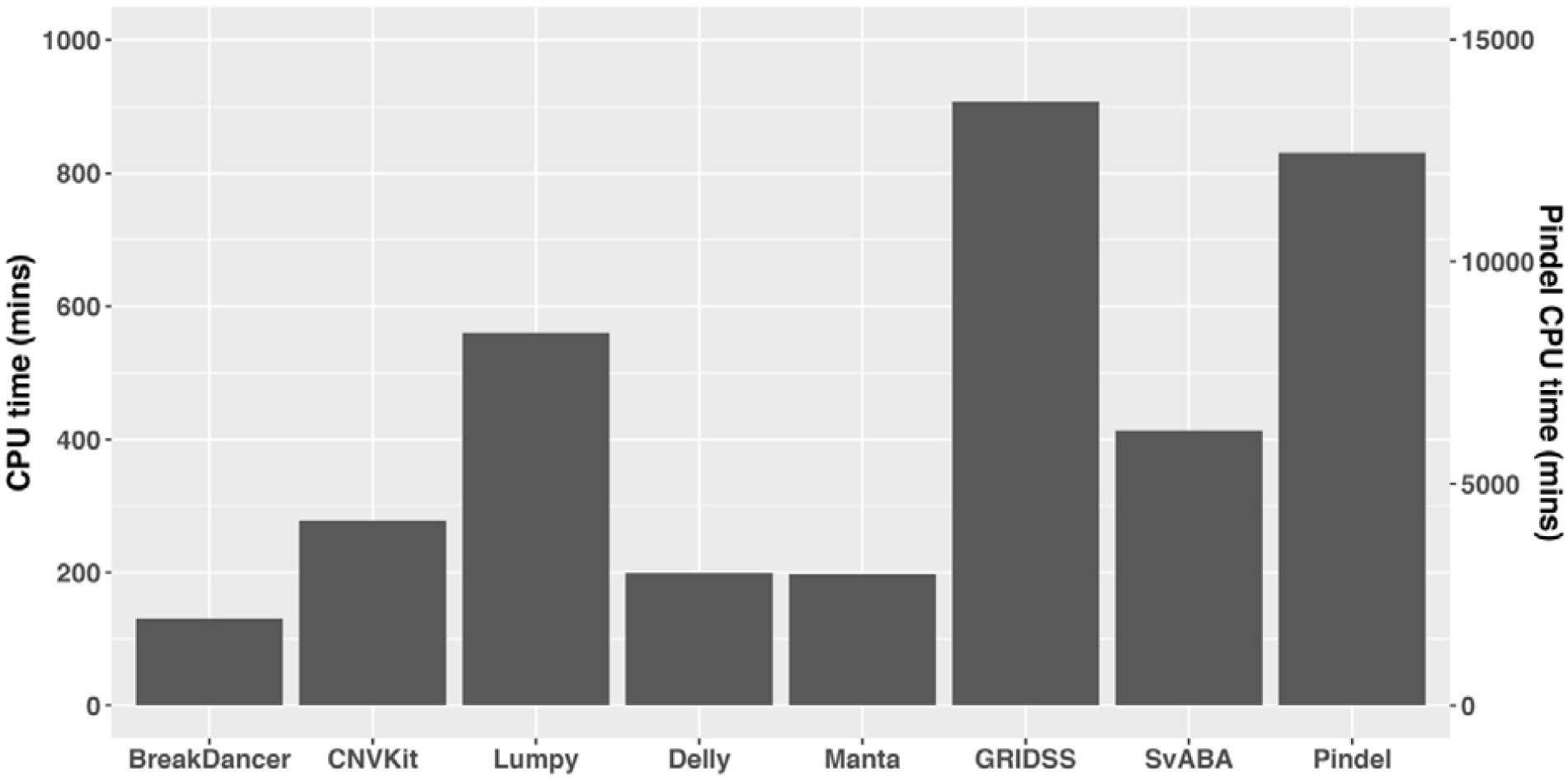
CPU time of structural variant callers. Results are based on simulation data with tumour and matched normal coverage of 60x and variant allele fraction of 100%. The CPU time of Pindel is labelled on the right vertical axis.

### Structural variant types and definitions

Broadly, SVs can be classified into five types (**Figure 4**). Deletion (DEL) is the removal of a DNA segment from the genome. Duplication (DUP), also known as tandem duplication, is the event of copying a DNA segment and inserting it beside the original copy. Inversion (INV) is the inversion of a DNA segment at the same locus. Insertion (INS) has two subtypes: domestic insertion (DINS) is the addition of a DNA segment copied from a distant site of the same genome (i.e. “copy-and-paste”) and foreign insertion (FINS) is the addition of a novel sequence, not known to be present in the sample genome. Translocation (TRA) involves the deletion of a DNA segment from one locus and its reinsertion at another locus (i.e. “cut-and-paste”). As a result, a translocation event is associated with a deletion event (DEL_TRA). Furthermore, in accordance with the Variant Call Format (VCF) 4.2 specification (updated 8 March 2019), a fusion junction in a rearranged genome can further be described by two adjacent breakends (BND) with coordinates relative to the reference genome, such that domestic insertions and translocations can be represented by two pairs of BNDs (**Figure 4**).

**Figure 4.**
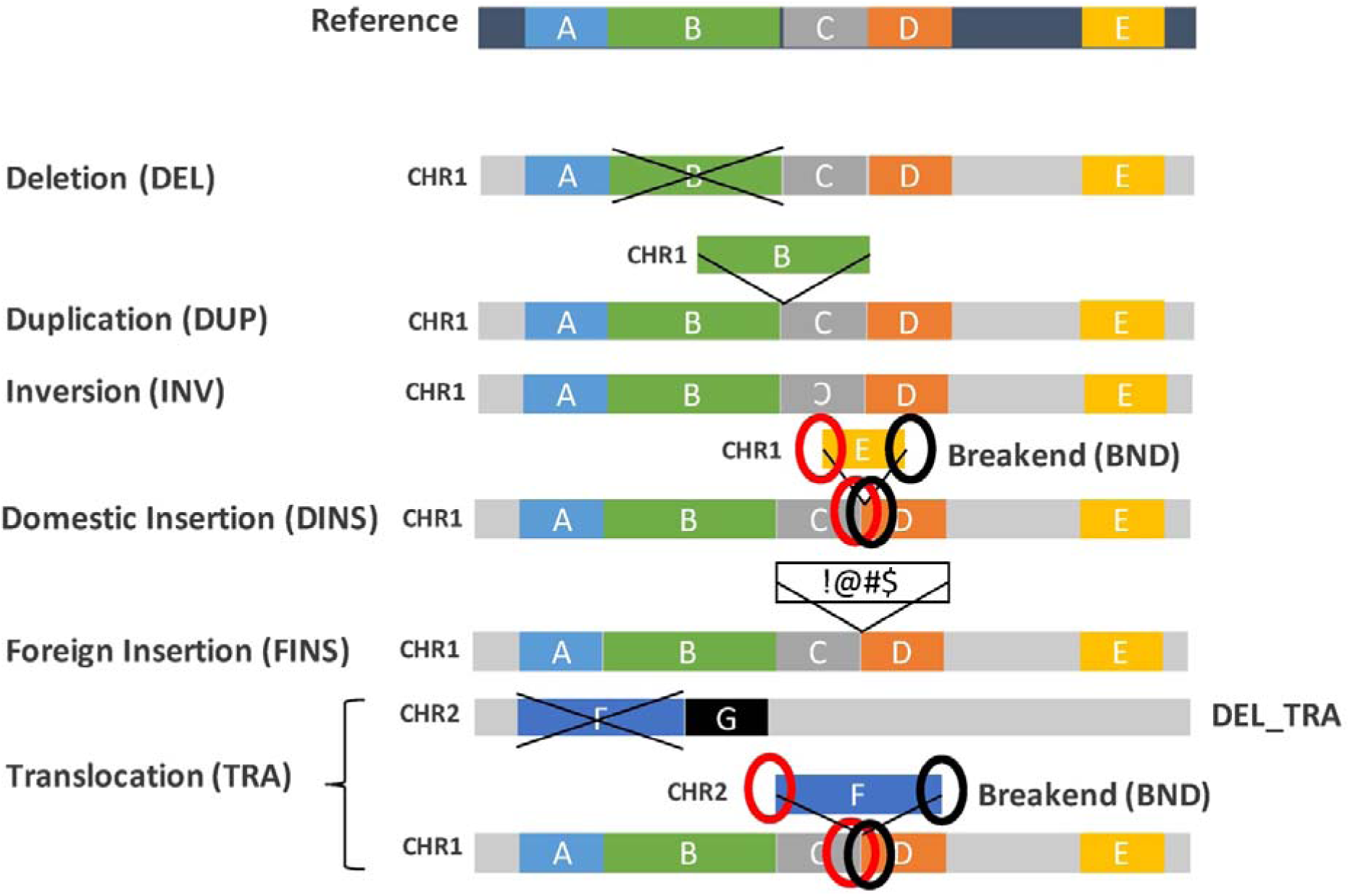
Definition of structural variant types. A “normal” representation of the reference genome (dark blue background) with five schematic regions (labelled A-E) is shown at the top. Below are different representations of rearranged genomes (grey background), relative to the reference, corresponding to each of the SV types: deletion (DEL), duplication (DUP), inversion (INV), insertions (domestic, DINS; or foreign, FINS) and translocation (TRA). DINS and TRA can be represented by two pairs of breakends (BND).

In spite of these broad classes, technical definitions for each SV category can vary between SV callers (**Table 2, Supplementary Methods**). BreakDancer can detect and reports DEL, INS, INV, intra-chromosomal translocation (ITX; which could also include DUP and DINS) and inter-chromosomal translocation (CTX; which could include DINS). CNVKit being a copy number variant caller, only reports copy number gains (DUP) and losses (DEL). Pindel reports DEL, DUP, INS, INV and replacement (RPL). A RPL describes an insertion event around the breakpoint of a deletion event, which could in fact capture DUP, INV, DINS, and FINS, thus occasionally resulting in duplicate SV calls with different assigned SV types. Lumpy and Delly both based on both read-pair and split-read alignment signatures can detect DEL, INV, DUP, and BND of inter-chromosomal events. Delly detects small INS that can be captured by single reads (smaller than read-length) whereas Lumpy cannot detect small INS. Neither can detect full intra/inter-chromosomal insertions with insert fragments exceeding the library insert size; evidence of (unpaired) breakpoints for these events are reported as BND. Manta can detect and report DEL, INV, INS (fully and partially assembled insertions), DUP, and BND of inter-chromosomal events. GRIDSS being an SV breakpoint caller initially reports all variants as BND but can be post-annotated as DEL, INS, INV, DUP and BND with its accompanying R script. SvABA, similar to GRIDSS, initially reports all SVs as BND with SV types further assigned, where possible, according to breakpoint orientations, as INV, DEL and DUP or INS (DUP/INS) which cannot be distinguished through breakpoint orientation.

**Table 2.**
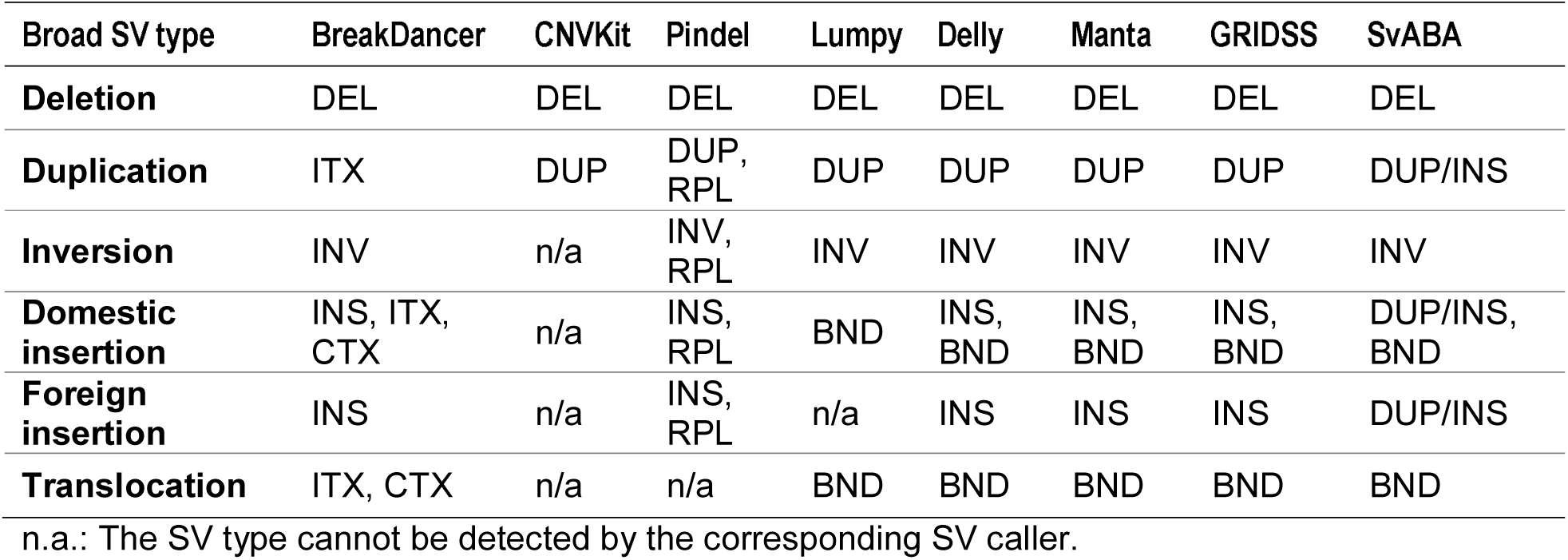
Comparison of structural variant types and definitions across SV callers.

| Broad SV type | BreakDancer | CNVKit | Pindel | Lumpy | Delly | Manta | GRIDSS | SvABA |
| --- | --- | --- | --- | --- | --- | --- | --- | --- |
| **Deletion** | DEL | DEL | DEL | DEL | DEL | DEL | DEL | DEL |
| **Duplication** | ITX | DUP | DUP,<br>RPL | DUP | DUP | DUP | DUP | DUP/INS |
| **Inversion** | INV | n/a | INV,<br>RPL | INV | INV | INV | INV | INV |
| **Domestic insertion** | INS, ITX, CTX | n/a | INS, RPL | BND | INS, BND | INS, BND | INS, BND | DUP/INS, BND |
| **Foreign insertion** | INS | n/a | INS, RPL | n/a | INS | INS | INS | DUP/INS |
| **Translocation** | ITX, CTX | n/a | n/a | BND | BND | BND | BND | BND |

n.a.: The SV type cannot be detected by the corresponding SV caller.

#### Deletions are the easiest to detect, while foreign insertions are the most difficult

Here, we evaluate each SV caller’s ability and efficiency to detect different SV types using a simulated somatic SV callset comprising of 200 of each of the six SV types, noting that this corresponds to a total of 400 DEL events with 200 of them due to translocation events (**Figure 5a**). BreakDancer reports a large number of ITX and CTX calls. While the majority (91%) of ITX calls are true DUP events (**Figure 5b orange**), most (94%) of the CTX are false positives (**Figure 5b green**). It performs modestly for DEL events with 73% recall rate but 52% FP rate. CNVKit performs poorly for both DEL and DUP, the only two SV types it is capable of detecting, with <2% recall rate and >96% FP rate. Interestingly, both BreakDancer and CNVkit have low sensitivity in detecting DEL_TRA (**Figure 5a**). This is because, unlike DEL, DEL_TRA events are not completely eliminated from the genome, but are moved to another genomic position. Consequently, there is no obvious reduction in depth of coverage for CNVKit to capture the deletions, and although BreakDancer can detect DEL_TRA, it does so with very poor breakpoint resolution (typically > 200 bp), reflecting the limitation of the read-pair based method as discussed. Pindel reported more than 28,000 somatic SV calls, of which 63% are INV and 34% are RPL calls. Considering an overall recovery rate of 45% (**Figure 2a**), Pindel exceeds this for two subclasses of SVs, recovering 69% DEL and 83% INV (**Figure 5a**).

**Figure 5.**
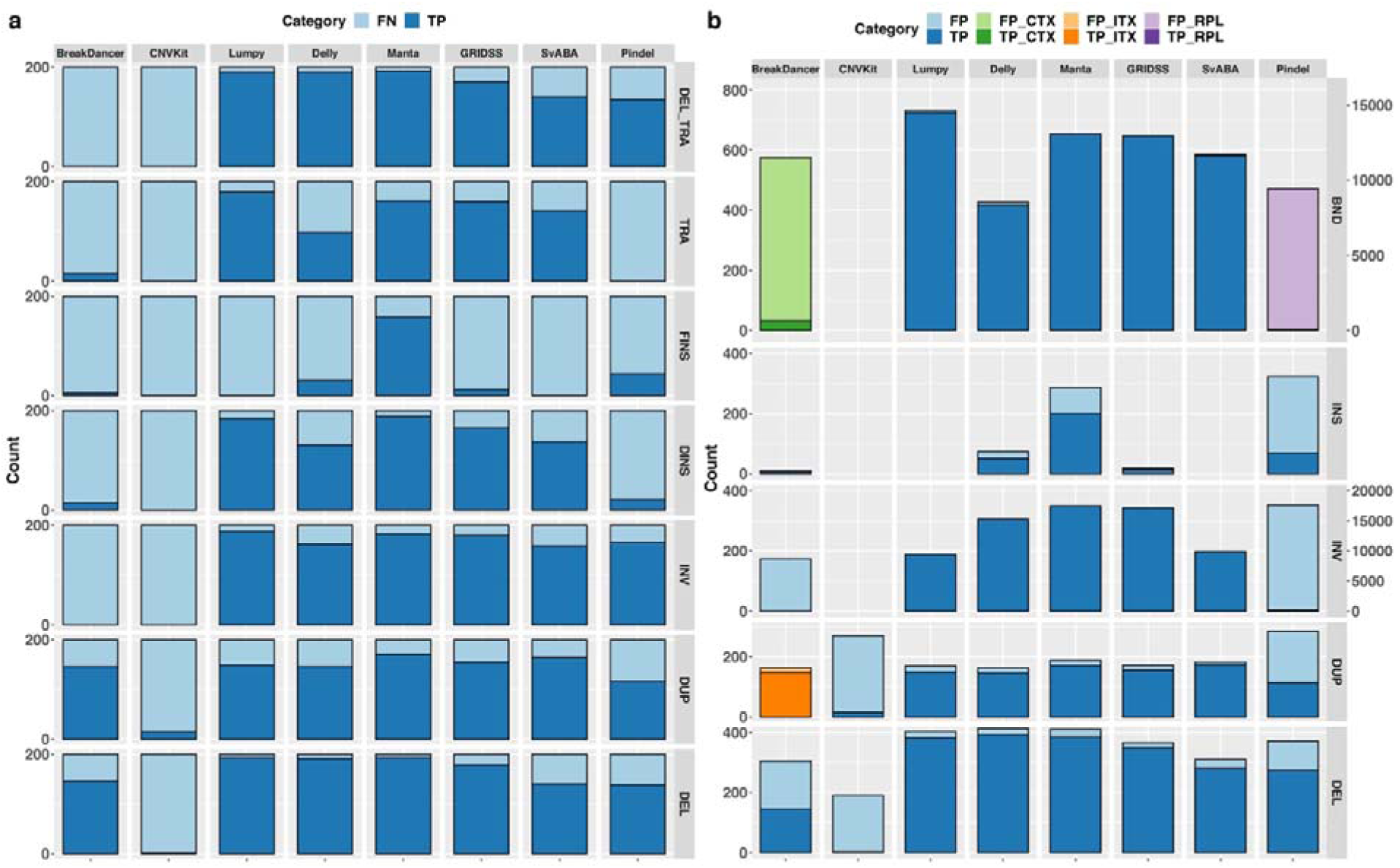
Performance of SV callers in detecting different SV types. (a) The number of simulated SVs recovered (dark blue; TP) or missed (light blue; FN) by individual SV callers for each SV category: deletion (DEL), duplication (DUP), inversion (INV), domestic insertion (DINS), foreign insertion (FINS), TRA (translocation), and TRA_DEL (deletion due to translocation). (b) The total number of SVs reported by individual SV callers that are true positives (dark shade; TP) or false positives (light shade; FP). The total number of DUPs shown under SvABA can be either DUP or INS events. The number of inter-chromosomal translocation (CTX) reported by BreakDancer are shown in the BND category in green. The number of intra-chromosomal translocation (ITX) reported by BreakDancer are shown in the DUP category in orange. The number of replacement (RPL) reported by Pindel are shown in the BND category in purple. The count of RPL and INV by Pindel is labelled on the right.

Overall, the other five SV callers perform well particularly for TRA and DINS events, with Lumpy and Manta achieving the highest sensitivity and missing only 9% and 6% of simulated SVs respectively (**Figure 5a**). Although these SV callers did not explicitly report TRA and DINS events, they can detect BNDs of rearranged fragments in inter-chromosomal events with high sensitivity and precision. Although Delly and Lumpy are both based on a combination of read-pair and split-read alignment signatures, Delly missed many more TRA and DINS (51% and 34%, respectively) than Lumpy. This is because Lumpy integrates both alignment signatures for SV discovery, while Delly uses split-read only for SV breakpoint refinement.

All SV callers struggled to detect simulated FINS, with 79% detected by Manta and <30% recovered by other SV callers (**Figure 5a**). Of the FINS recovered by the SV callers with the exception of Manta, all were < 500 bp, which is the mean insert size simulated. This poor performance in FINS detection is from all callers being reliant on the initial alignment of sequencing reads to the reference genome. In case of FINS, where the inserted sequence is absent in the reference genome, the corresponding reads are typically missed early in the SV detection pipeline, or even filtered out prior to SV detection. Therefore, FINS can only be inferred from reads that align in a split-read manner or reads whose mates are properly aligned, resulting in only small FINS within the library insert size range can be detected. Differently, Manta report large insertions even though the inserted sequence cannot be fully assembled, resulting in higher recall rate in FINS detection. However, Manta had 30% false positive INS calls, significantly higher than other SV callers (**Figure 5b**). Interestingly, those false positive INS calls were found in small sizes (< 102 bp) and reported to wrong SV types.

It should be noted that, while Manta, GRIDSS and Delly reported more inversion events than were simulated (**Figure 5b**), they do not have a proportionately higher false positive rate. This is because these three SV callers report two paired-end clusters for each INV (INV3 and INV5 tags for Manta and CT = 3to3 or 5to5 tags for Delly in VCF/INFO field and 2 pairs of BND for GRIDSS). Taking this into consideration, the three SV callers are able to recover >80% of INV while missing <10% (**Figure 5a**).

In summary, SV callers based on more than one method in the discovery step have better overall performance across all SV types, except for FINS. As expected, DEL is the “easiest” to recover. Interestingly, INV and TRA have similar recovery rates as DUP, suggesting the relatively fewer reported cases of INV & TRA in the literature is due to these two SV types being genuinely less prevalent in cancer genomes than DUPs, rather than it being a methodological limitation. Among the eight SV callers examined, Manta and Lumpy performed the best, with most notable differences in Manta being able to detect DUP better (7.6% higher F1 score) but Lumpy being able to detect BND better (5% higher F1 score; **Supplementary Table S1**).

### Impact of structural variant sizes on SV detection

SVs are distinguished from small indels by the number of bases they impact with 50 bp being the commonly accepted threshold for classification as SVs. Here, we compared the SV callers’ ability to detect SVs at different size ranges, from 50 bp to 1 Mbp. Most SV callers are consistent in the overall number of SVs detectable across different size ranges (**Figure 6**). CNVkit is the exception, where the extent of reported SVs appears to increase with SV size. We note here that, BreakDancer is unable to estimate the size of translocated DNA fragments and so assigns an arbitrary 498 bp for all reported CTX events; these are excluded in **Figure 6**.

**Figure 6.**
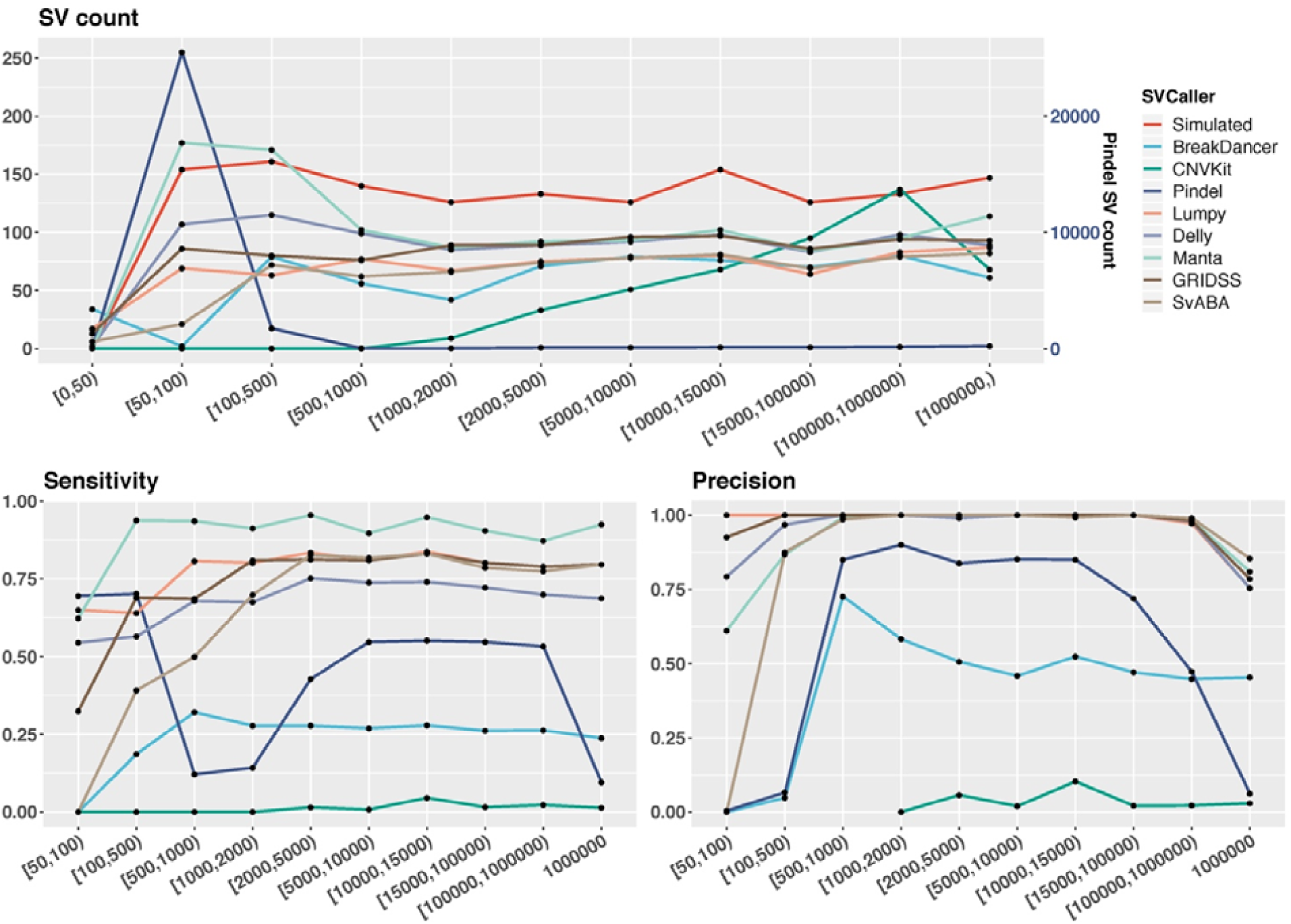
Performance of structural variant callers across different variant size ranges. Shown on top are the number of SVs reported by each SV caller within each defined SV size range. The bottom plots show sensitivity (left) and precision (right) evaluated based on a breakpoint precision threshold of 200 bp. Results are based on simulated samples with tumour and matched normal coverage of 60x and tumour purity of 100%. Size of TRA correspond to the size of the translocation DNA fragment. The Pindel SV count is on the right vertical axis.

All SV callers struggled to detect SVs < 100 bp, with Pindel achieving highest sensitivity of 70%, which is still lower than the highest sensitivity for other size ranges. CNVkit is unable to detect SV < 1,000 bp due to the segment (window) size of 842 bp used, which was estimated by the caller (**Supplementary Methods**). In general, most SV callers detect small SVs (<100 bp) with lower precision, with GRIDSS and Lumpy being the exceptions (> 90%). Additionally, all SV callers have lower precision in detecting large SVs (1 Mbp), with SvABA having the highest precision at 85%. However, SvABA has zero sensitivity and precision in detecting SVs within 50-100 bp. In fact, the smallest SV size reported by SvABA is 97 bp; this is due to an arbitrary cutoff used by SvABA (taken from BWA-MEM) to distinguish SVs and indels. This likely explains its overall lower performance compared to other multi-method SV callers.

In conclusion, SV callers based on more than one SV detection methods show highest consistency in performance across the different SV size ranges. However, there are still limitations to accurately detect SVs >1 Mbp for all SV callers. The split-read based method shows its power in detecting smaller SVs (50-100 bp), though it needs to be well incorporated with other methods to attain a good performance in both sensitivity and precision.

### Breakpoint precision of structural variant calls

Breakpoint features are important for elucidating the underlying mechanisms of SVs, such as micro-homology-mediated DNA repair and movements of transposable elements [24]. Precise SV breakpoint positions can also facilitate accurate annotation of SVs, including impact on transcript splice sites and regulatory elements [17]. Furthermore, when comparing SV callsets, there is a need to define sufficient closeness between two calls for consideration as identical calls [10]. To better understand the extent of breakpoint resolution across the SV callers, we evaluate the change in the number of true positive calls at varying breakpoint precision threshold from 0 to 2,000 bp (**Supplementary Methods**). Breakpoint resolution is defined as the absolute distance between a true (simulated) breakpoint position and that reported by an SV caller (d_1_ and d_2_ in **Figure 7**).

**Figure 7.**
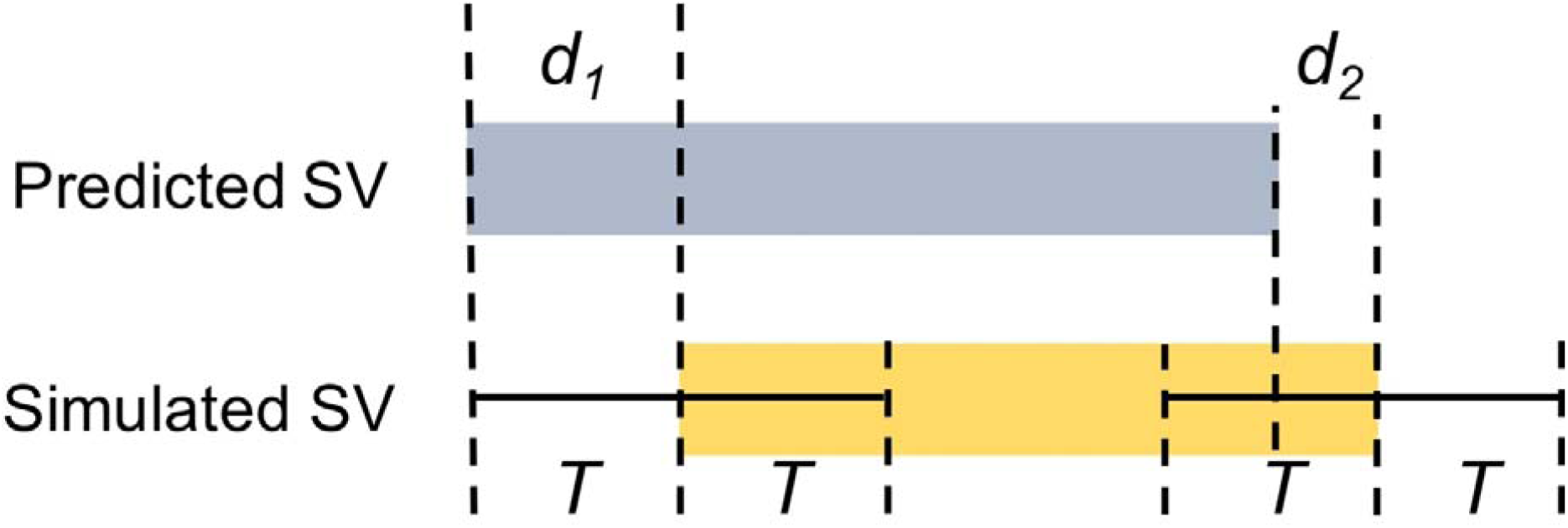
Breakpoint resolution threshold definition. A breakpoint resolution threshold, T, is defined as the absolute distance from a simulated breakpoint position within which a reported SV must fall. Thus, in this diagram, both d1 and d2 must be less than the defined threshold, T, for the predicted SV to be considered a true positive call.

#### SV callers using split-read method consistently detect structural variants within 2 bp resolution

BreakDancer and CNVKit have very poor breakpoint resolution (**Figure 8**). BreakDancer requires read-pairs to completely span the breakpoints and lie within either side of an SV event. Therefore, breakpoint locations can only be approximated by the mapping position of discordant read-pairs; actual breakpoints are not captured. Poor breakpoint resolution in CNVKit is due to the read-depth method, estimating average sequencing coverage within a pre-defined genomic interval (or bin). So, as long as the majority of an SV is within a genomic bin, there will be sufficient signal for a CNV call; the location of the breakpoints is not captured. Pindel has great breakpoint precision with 95% of all SV calls within 2 bp resolution, reflecting the advantage of using split-read signatures. Again, SV callers using multiple SV detection methods have better overall breakpoint resolution, with 99% of breakpoints called within 2bp for Manta and GRIDSS, within 5bp for Lumpy and within 100 bp for SvABA.

**Figure 8.**
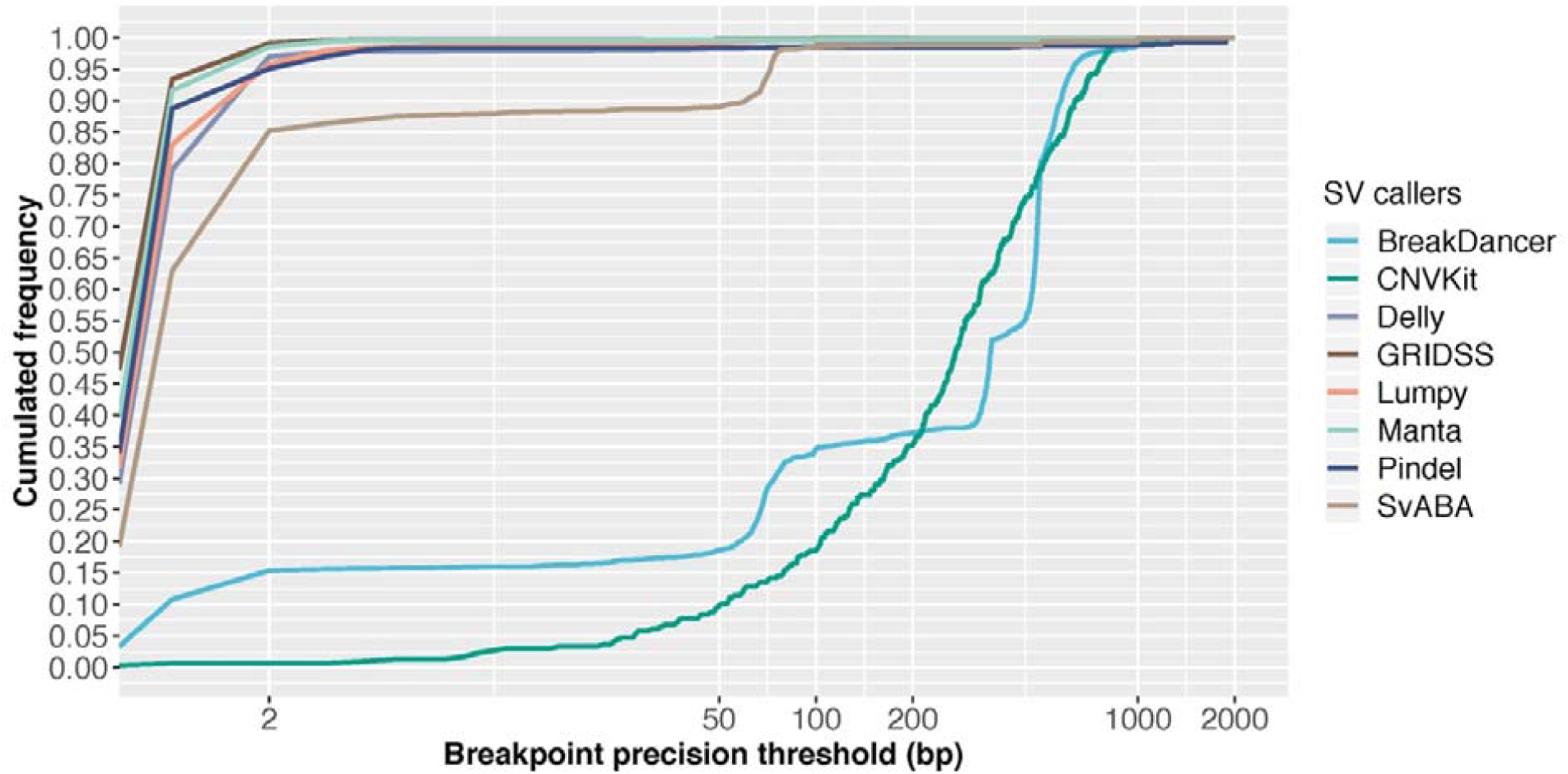
Structural variant calling sensitivity at increasing breakpoint resolution threshold. Results are based on simulation data with tumour and matched normal coverage of 60x and variant allele fraction of 100%. This figure shows the cumulated frequency of the number true positive structural variants detected by each caller for breakpoint precision thresholds less than 2,000 bp.

While several SV callers are able to achieve near perfect breakpoint resolution (within 2 bp), we observed a loss in precision with each additional processing step for all SV calling pipelines. In particular, we noted that between SV calling and SV annotation, reported breakpoints fluctuate by one or two bp, largely due to conversions between 0-based and 1-based coordinates required for the different file formats (namely, BAM to VCF to BED and sometime back to VCF).

In addition, micro-homology around SV breakpoints can confound sequence alignment, which also contributes to imprecision of breakpoint prediction. To be mindful of micro-homology, users can check the confidence interval for breakpoint positions (CIPOS and CIEND) and base pair identical micro-homology length and sequence (HOMLEN and HOMSEQ) in the output VCF files, as specified in the VCF 4.2 specification (updated 8 March 2019). However, of the eight SV callers examined in this study, only GRIDSS and Manta report all of these, while Lumpy only reports CIPOS and CIEND and SvABA only provides HOMOLEN and HOMSEQ. In summary, SV callers based on more than one SV detection method have higher breakpoint detection resolution and split-read method is essential for precise breakpoint detection.

### Impact of variant allele frequency on structural variant detection

Variant allele frequency, or variant allele fraction, (VAF) is the proportion of sequencing reads supporting the detected variant at a given locus. In cancer genomics, two key factors influence VAF: tumour purity and tumour heterogeneity. Tumour purity is the proportion of cancerous cells captured in a sample, which is typically uncontrollable without the support of imaging-guided biopsy or histopathology-guided dissection. Tumour heterogeneity refers to the molecular diversity between cancer cells of a tumour sample and reflects the amount of genomic change throughput the development of the cancer. Obviously, SVs at low VAF are more difficult to detect than SVs in high abundance. However, detection of SV at low VAF can theoretically be compensated by increasing sequencing coverage [7]. In this review, we provide a comprehensive evaluation of the relationship and joint effect of VAF and sequencing coverage on SV detection from five SV callers, across different SV types and for different SV sizes. BreakDancer, CNVKit and Pindel were excluded in this analysis due to their notably poor performance even with 100% VAF as shown above (**Figure 2**) and previously by others [8].

#### Variant allele frequency has a non-linear impact on sensitivity and little impact on precision

Overall, VAF has a monotonic but non-linear effect on sensitivity (**Figure 9a, 9c**), and little impact on precision (**Figure 9b, 9d**). The positive effect of VAF on sensitivity is greatest for lower VAF, with signs of saturation at VAF > 0.2 for all SV callers except Delly. While Delly appears to perform similarly to other SV callers at high VAF (> 0.5), its performance is substantially worse at low VAF dropping to zero sensitivity at VAF < 0.1. In addition, at 0.1 VAF, Manta, Lumpy and GRIDSS maintain overall sensitivity at around 60%, whereas SvABA sensitivity drops to 30%.

**Figure 9.**
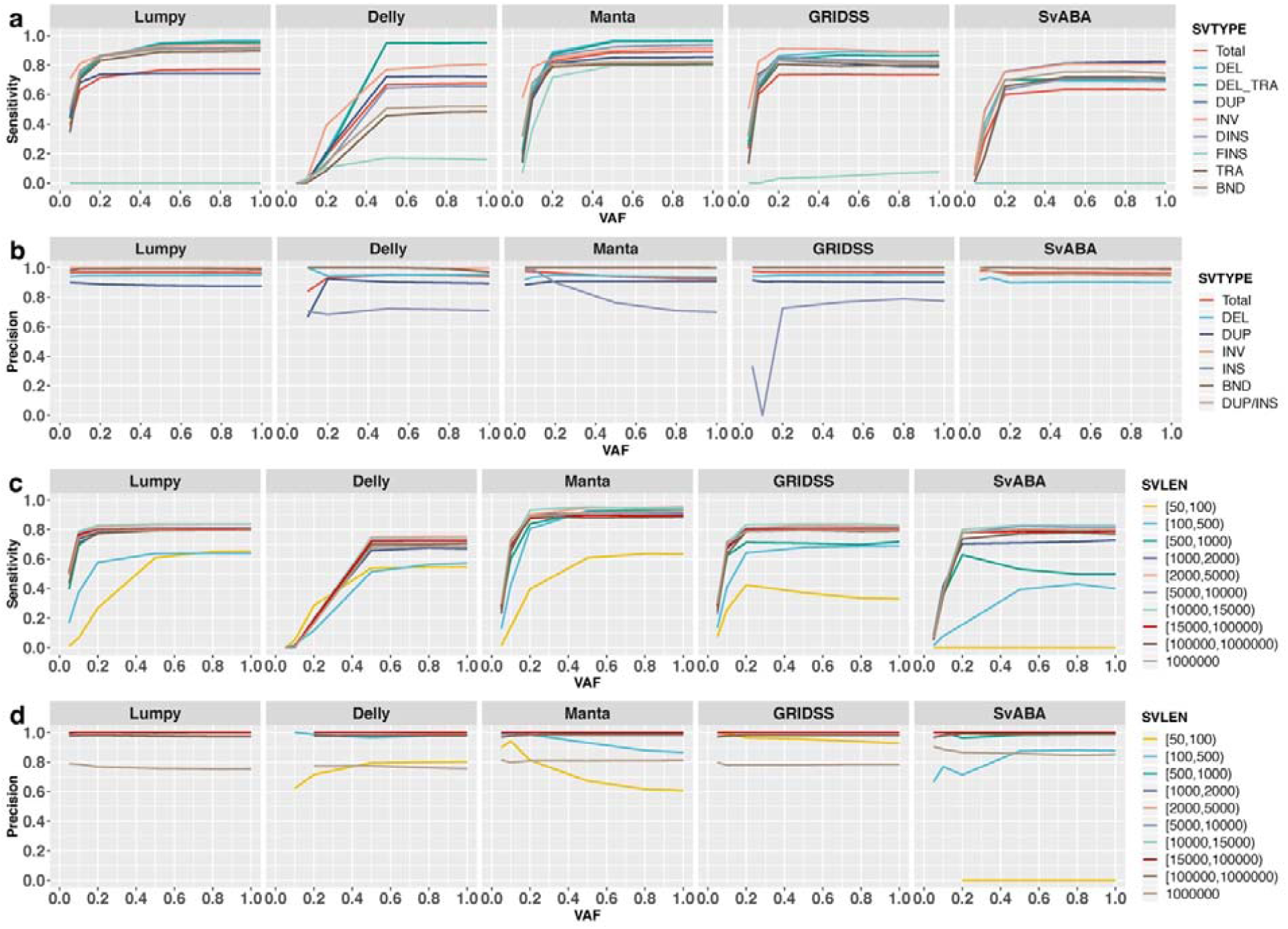
Impact of variant allele frequency on the detection of different SV types and sizes. Shown are sensitivities (a and c) and precisions (b and d) for different SV types (a and b) and SV size ranges (c and d) using five SV callers: Lumpy, Delly, Manta, GRIDSS, and SvABA. Results are based on simulation data with tumour and matched normal coverage of 60x, and breakpoints within 200 bp of simulated SVs.

This overall effect of VAF on SV detection sensitivity is similar for different SV types, with one exception: VAF has little to no effect on FINS detection. This is unsurprising as FINS is difficult to detect for all SV callers even at 100% VAF (**Figure 5**). Interestingly, increasing VAF appears to negatively impact the precision of INS detection for Manta. This is due to the rapid increase in the absolute number of INS called by Manta with increasing VAF (10 INS at VAF = 0.05, 66 INS at VAF = 0.1 and 171 INS at VAF = 0.2), of which, many are false positives. The fluctuation in precision for INS detected by GRIDSS at varying VAF (**Figure 9b**) is due to the low numbers of total INS detected (only one INS at VAF ≤ 0.1, 12 INS at VAF = 0.2, and 22 INS at VAF = 1). Again, Delly behaves differently compared to other SV callers in that, there is more distinct variability in its performance for different SV types. In particular, Delly performs much better for DEL than other SV types, especially at VAF > 0.5.

As previously observed (**Figure 6)**, small SVs (50-100bp) and large SVs (> 1Mbp) are more difficult to detect. However, the impact of VAF to detect different SV sizes is similar across the SV callers. The most notable difference is the counterintuitive decrease in precision for Manta for detecting SV < 100 bp with increasing VAF (**Figure 9d, Manta**). Many of the false positive SVs < 100 bp are INS events, corroborating with the similar trend observed above (**Figure 9b, Manta**), suggesting Manta has elevated false positive calls for small INS especially with increasing VAF. Furthermore, many of the false positive small INS appear to be a result of incorrect SV type assignment by Manta; i.e. DUP, INV and BND of TRA events reported as INS.

#### Deep sequencing is critical when variant allele frequency is less than 50%

To determine if, and to what extent, increasing sequencing depth can improve SV detection at low VAF, we evaluated six levels of tumour (20x, 30x, 45x, 60x, 75x and 90x) and normal coverage (15x, 30x, 45x, 60x, 75x and 90x) at six VAF levels. As expected, increasing tumour sequencing coverage can greatly improve somatic SV detection sensitivity for samples with low VAF (**Figure 10a and 10b**). For example, at 0.2 VAF, increasing tumour coverage from 20x to 30x can increase sensitivity by 20% using Manta. However, the benefit of increasing coverage quickly saturates. For example, at 0.2 VAF, increasing tumour coverage from 60x to 90x yields only a 5% improvement in sensitivity for Manta. In contrast, the impact of tumour sequencing coverage has less impact for samples with higher VAF. For example, at 0.5 VAF, Manta sensitivity increases only by 6% from 20x to 30x and 1% from 60x to 90x. The absolute extent of impact from tumour sequencing coverage is different for different SV callers. In particular, Delly gains < 10% sensitivity from 20x to 30x depth of tumour coverage and derives no further gain above 30x. It is notable that tumour coverage has almost no impact on the precision of SV detection regardless of VAF level (**Figure 10b**). The small fluctuation on Delly’s precision at low VAF is due to the small number of total detected SVs.

**Figure 10.**
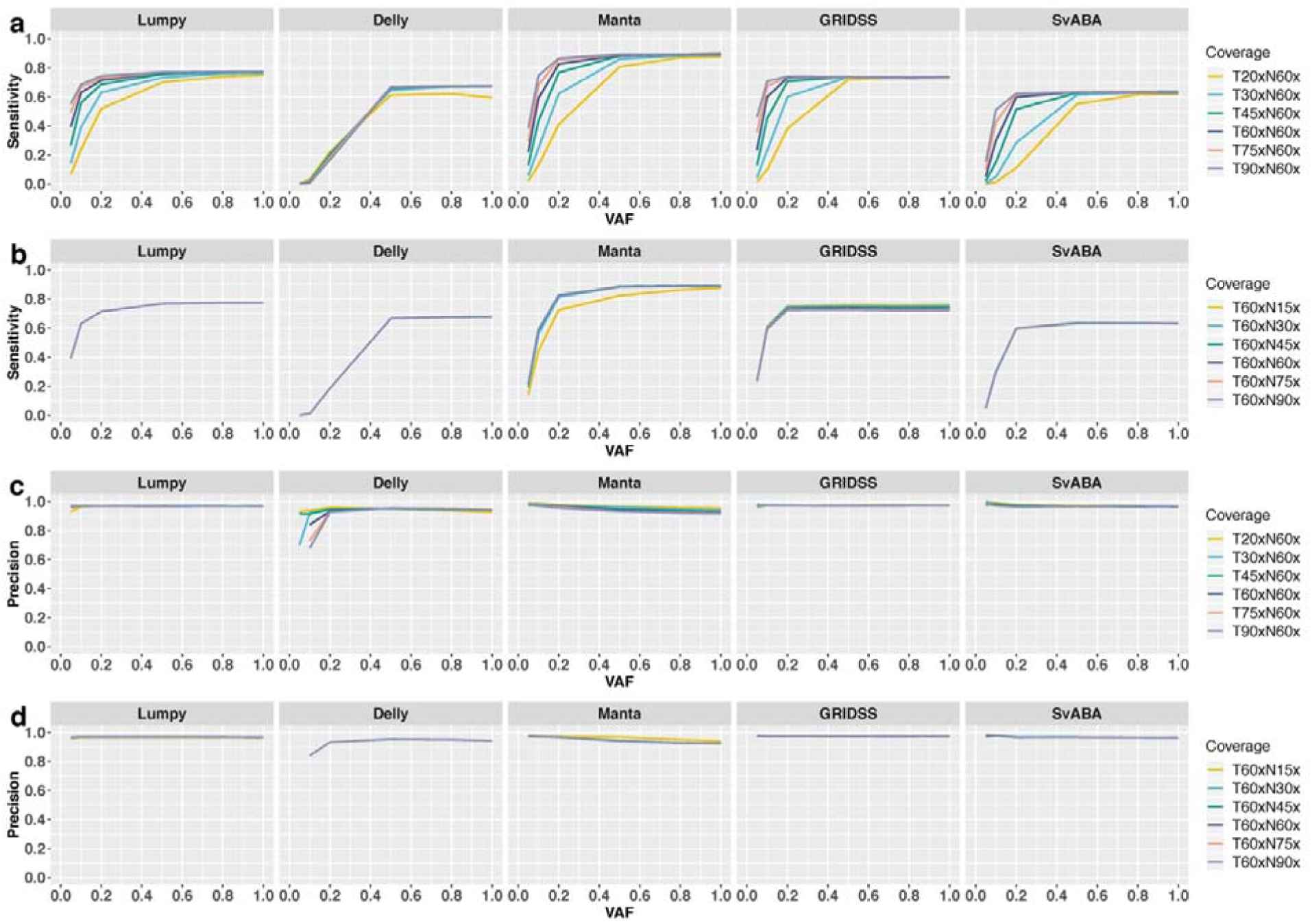
Joint impact of sequencing coverage and variant allele frequencies on structural variant detection. Shown are total sensitivity (a) and precision (b) across variant allele frequencies (VAF) with different sequencing coverage of the tumour samples (a and c) and match-normal samples (b and d), based on breakpoint precision threshold of 200 bp.

In contrast to tumour sequencing depth, sequencing coverage of the matched-normal sample has little impact on either sensitivity or precision, with the exception that increasing normal-sample coverage from 15x to 30x improves sensitivity for Manta (**Figure 10c**).

In summary, when VAF is high (> 0.8), little value can be gained from increasing sequencing depth. In fact, when VAF is close to 1, all evaluated SV callers have reasonable sensitivity (>60%) to identify somatic SVs at low sequencing coverage (20x-30x). In contrast, deep tumour coverage (75x-90x) is critical when VAF is low (< 0.5). Finally, when VAF is less than 0.1, there is little to no power in SV discovery even for deeply sequenced (>90x) tumour samples.

### Impact of segmental duplication on structural variant detection

Genomic regions of low complexity, such as repeats and GC-rich regions [2], can result in ambiguous read-mapping, which can lead to incorrect read-alignments and subsequent false variant detection. SV callers usually use a mapping quality threshold to ensure SVs are supported by unique mapping. Ambiguous mapping can also affect precise breakpoint prediction. In particular, segmental duplications (SegDup), which comprise around 5% of the human genome, are sequences at different genomic loci that share a high level of (>90%) sequence identity [25], and is one of the biggest sources of false variant calls. In a recent study, it was found that germline SVs at low complexity and simple or short tandem repeat regions have lower precision at different levels for all callers [9]. Here, we evaluate the impact of segmental duplication on somatic SV calling.

#### Segmental duplicated regions are prone to false positive duplication and deletion calls

On average, somatic SV residing within SegDups are 2.2% (0.7%-12%) less likely to be detected and induce 14% (11%-18%) higher false positive calls than those outside of SegDup regions (**Figure 11**). The most significant impact on detection sensitivity was observed for FINS called by Manta and DELs from Delly. Unlike VAF and sequencing coverage, which affect sensitivity more than precision, the reverse is true for SegDups. Across all SV callers, precision is greatly reduced for SV collocating at SegDup regions, especially for DUP and DEL events; that is, false positive DUP and DEL are mainly due to SegDup regions.

**Figure 11.**
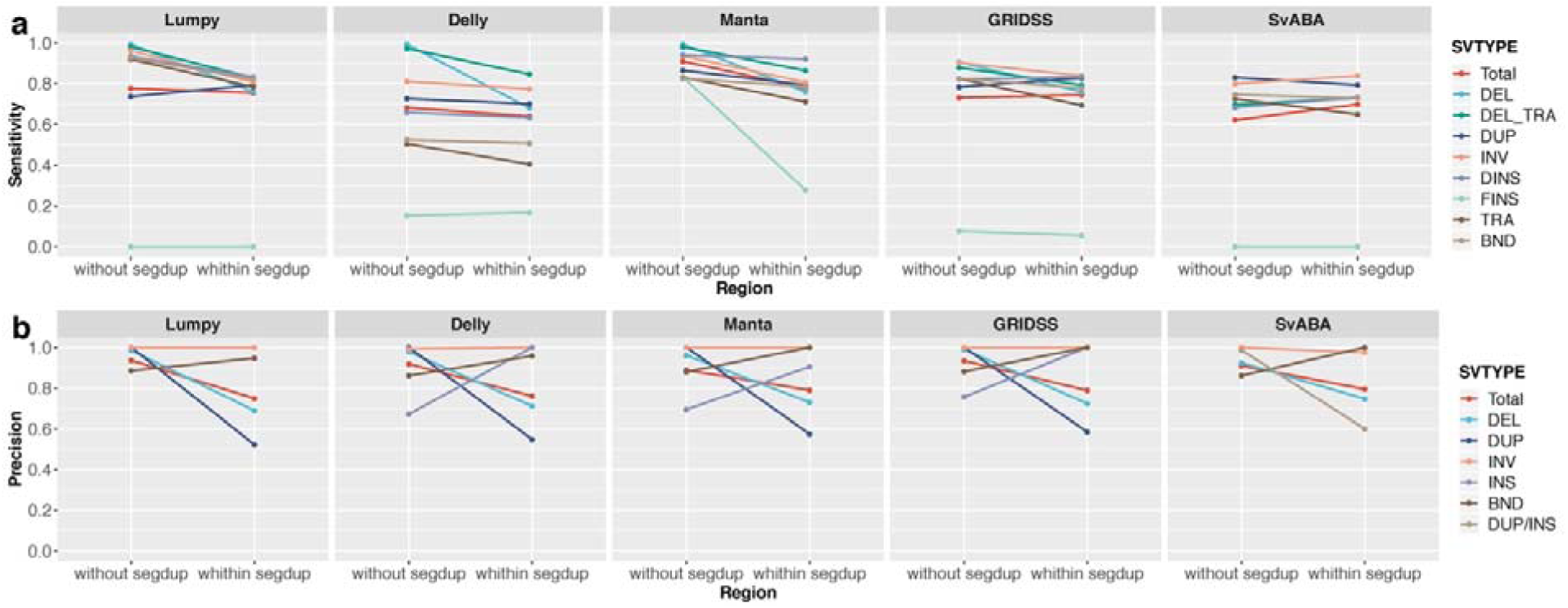
Sensitivity and precision of structural variant detection within and outside of segmental duplicated regions. Results are based on simulation data with tumour and matched normal coverage of 60x, variant allele fraction of 100% and breakpoints within 200 bp of simulated SVs.

## Concluding remarks

Structural variants are an important type of genomic alterations in cancer, but are intrinsically more difficult to detect than small variants from short-read NGS data. While the surge of new SV callers in the past few years have significantly enhanced our ability to detect SVs from genomics data, it has resulted in the unintended effect of “overchoice”. To overcome this, recent studies have attempted to compare the performance of a variety of SV callers, but these have focused predominantly on germline SVs and simple SV types [8,9] and only on overall performance for somatic SVs [10]. In this review, we have added to this mounting evaluation effort by examining the effect of major factors affecting the ability of different methods in detecting somatic SVs. Although simulated data does not truly reflect the complexity of real cancer samples, the evaluation presented in this review provides an insight into the upper bound of what is achievable with current methodologies [9,10].

Overall, we recapitulate previous observations that SV caller based on more than one method performs better than those relying on single methods [8,9]. Furthermore, while SV callers based on the same detection method(s) have similar overall performance, they vary in their ability to detect different SV types and SV sizes. Therefore, for comprehensive detection of somatic SVs, it may still be necessary to use a combination of callers. From our evaluation, we found the pairwise union callsets of Manta and Lumpy or GRIDSS to provide the highest F1 value.

Moreover, we found that DELs tend to have a higher validation rate than other SV types. This likely explains the higher validation rate for DEL events [17]. However, we note that all SV callers have low discovery rate in calling novel sequence insertion (foreign insertion) and relatively small SVs (<100 bp) and have high false discovery rate in calling large SVs (> 1Mbp) and SVs with highly homologous breakpoints. This is an area for improvement, either for future somatic SV callers for NGS data or for long-range sequencing technologies. For factors affecting SV detection, we have found VAF, due to tumour purity and/or intra-tumour heterogeneity, to have the biggest impact. As expected, sequencing coverage of the tumour sample can rescue some of the sensitivity lost, however, the amount of improvement is not linear and differs between SV callers. In general, little improvement is observed (< 4%) beyond 60x depth of tumour coverage for VAF > 0.2 and there does not appear to be a significant impact from sequencing depth of coverage of the match-normal sample.

Additionally, and uniquely to this study, we evaluated SV callers’ ability in identifying inter-chromosomal rearrangements, such as DINS (“copy-and-paste”) and TRA (“cut-and-paste”) with inserted sequence from another chromosome. Due to inherent limitations of NGS data, inserted or translocated sequences cannot be fully captured. However, by jointly utilising read-pairs, split-reads and local-assembly methods, breakends of rearranged genomic fragments can be accurately resolved. We have shown that BND pairs of DINS and TRA are reported with high sensitivity and precision by SV callers that jointly use these methods (namely, Manta, GRIDSS, and SvABA). Correct and precise identification of BNDs are important as they are essential for SV annotation.

As the detection of small variants become routine and embedded within cancer genomics pipelines, it is imperative to start to consider and work towards capturing one of the most important genomic aberrations in cancer. As SV callers begin to saturate the bioinformatics field, it is timely to evaluate areas ripe for improvement. This review highlights these areas being the identification of foreign insertions and detection of relatively small and very large SVs. Moreover, there is still the unattained goal of an SV caller capable of identifying all SV types and sizes and capable of capturing the full inserted or translocated sequences.

## Key points

- A comparison of eight most commonly used structural variant callers for cancer genomics was performed
- Structural variant callers based on multiple detection methods are more sensitive and reproducible
- Addition of a second SV caller to an already high-performing caller adds little value
- Most SV callers have approximately 2 bp breakpoint resolution
- Variant allele frequency has a logarithmic impact on sensitivity but little effect on precision

## Competing interests

The authors declare that they have no competing interests.

## Funding

This work was supported in part by a Movember Revolutionary Team Award funded by Movember Australia and Prostate Cancer Foundation Australia. TG is supported by an Australian Government Research Training Program Scholarship. VMH is supported by the University of Sydney Foundation and Petre Foundation, Australia.

## Acknowledgements

We acknowledge the high-performance computing resources generously provided by the National Computational Infrastructure (Raijin), the Garvan Institute of Medical Research (Wolfpack) and the University of Sydney (Artemis).

