## Supplementary Methods for "Detection of somatic structural variants from short-read next-generation sequencing data"

### Somatic SV simulation framework

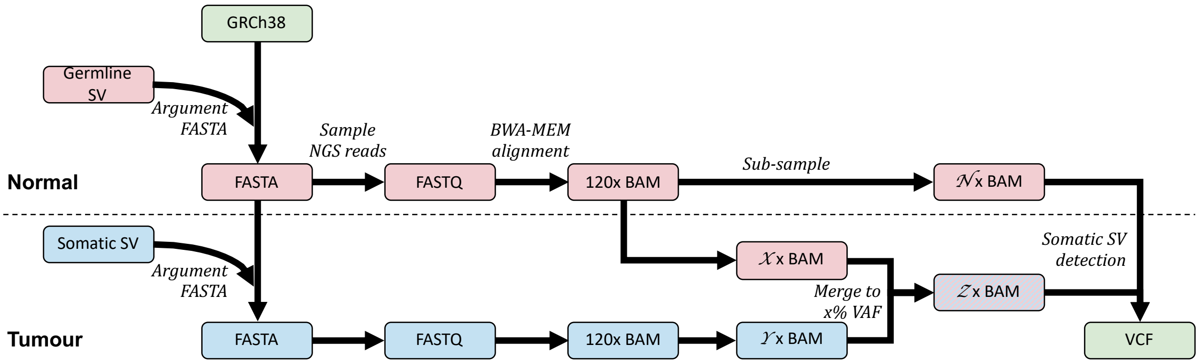


**Figure S1. Simulation framework.** A flowchart showing the simulation process. SVEngine is used to simulate tumour and normal genomes (FASTA) with somatic and germline SVs (BED), generating short-read sequences (FASTQ). Simulated short reads are aligned to the human reference genome GRCh38 using BWA-MEM to generate matching tumour and normal (germline) BAM files. Different depths of coverage are simulated by subsampling from the original bam files. Tumour purity is simulated by subsampling and merging different ratios of normal and tumour reads from the respective BAM files, to generate the final tumour BAM files. Somatic SV detection is performed for each tumour and normal pair and evaluated by comparing to the simulated somatic SV set.

##### Simulating genomes

Normal and tumour genomes were simulated based on the human reference genome GRCh38, to which we added a set of baseline SVs to emulate the germline genome, using SVEngine [11]. Duplicating this genome (fasta), we further added another set of SVs to emulate the cancer genome (**Figure S1**). In both cases, simulated SVs were distributed randomly along the genome, masking gap, centromeric and telomeric regions [11,26], with telomeres defined as 10 Mbp at both ends of each chromosome. Gap and centromeric regions were based on *gap* [last updated 24/12/2013] and *centromeres* [last update 17/08/2014] tables from the University of California at Santa Cruz (UCSC) Genome Browser. No SV fragment or breakpoint are overlapping.

Each of the two sets of SVs included 200 deletions (DEL), 200 duplications (DUP), 200 inversions (INV), 200 domestic insertions (DINS), 200 foreign insertions (FINS) and 200 translocations (TRA). Foreign (novel) sequences for FINS were simulated using the Bioconductor package *RBioinf* [27] without any constraints (i.e. each nucleotide was independently sampled from the set of {A,C,G,T} with equal probability). For each SV class, variable lengths were simulated with each containing SV sizes of 50 bp, 100 bp, 500 bp, 1,000 bp, 2,000 bp, 5,000 bp, 10,000 bp, 15,000 bp, 100,000 bp and 1,000,000 bp. Note that the SV length of a translocation refers to the length of the translocated piece of DNA fragment.

#### Simulating short-read sequences

Paired short-read sequences (fastq) were sampled from the simulated germline and tumour genomes using SVEngine (Xia et al.,2018) to 120x depth of coverage. Insert size of read-pairs was simulated with a normal distribution with mean of 500 bp and standard deviation of 100 bp, while read length was set to 150 bp. Other parameters were kept as SVEngine default. The two fastq files were each aligned to GRCh38 using BWA-MEM v0.7.17-r1194 [12] to generate matching tumour and normal bam files.

Different depth of coverage was simulated by subsampling from the two 120x dataset using Picard DownsampleSam (http://picard.sourceforge.net). Six depths of coverage (90x, 75x, 60x, 45x, 30x and 20x) for the tumour sample and six (90x, 75x, 60x, 45x, 30x and 15x) for the normal sample were assessed. Tumour purity was emulated by subsampling then merging different ratios of normal and tumour reads from the respective bam files, using Picard MergeSamFiles, to generate the final tumour bam files. For example, tumour purity of 50% for a 60x tumour sample would be generated by merging an equivalent of 30x normal reads and 30x tumour reads from the respective bam files. Six tumour purity values (100%, 80%, 50%, 20%, 10% and 5%) were included for evaluation. It is worth noting that, as the detection of each SV is independent of each other, each simulated tumour purity (in reference to a sample) is effectively the simulated variant allele frequency (VAF) of each SV given only a single cancerous population in the corresponding dataset.

In all, 216 pairs of BAM files corresponding to all permutation of six depths of coverage of the normal samples (15x, 30x, 45x, 60x, 75x, 90x), six depths of coverage of the tumour samples (20x, 30x, 45x, 60x, 75x, 90x) and six VAF (5%, 10%, 20%, 50%, 80%, 100%) were generated. To account for stochastic noise, each of the 216 in silico experiments were replicated three times for a total of 648 datasets. The averaged results across the three replicates are presented in the paper.

#### Somatic SV calling

Somatic SVs were called using eight SV callers: Manta (v1.4.0), Lumpy (v0.2.13), GRIDSS (v 1.8.1), BreakDancer (v 1.4.5), Pindel (0.2.5b9), CNVKit (0.9.4), SvABA (v0.2.1) and Delly (v0.7.8) for each tumour/normal pair. BreakDancer, Pindel and CNVKit were performed on only one simulated experimental design, namely tumour and normal coverage at 60x and VAF of 100%. The remaining five SV callers were run on all 648 simulated datasets.

The commands and options used for each SV callers in this study are described below. For all SV callers, the input data was a pair of tumour-normal alignment BAM files, denoted here as TUMOR_BAM and NORMAL_BAM respectively. For some SV callers, the procedures for further filtering on high-confidence somatic SVs are explained.

##### BreakDancer

A configuration file was first created with bam2cfg.pl, filtering on minimum mapping quality PHRED score of 15 (-q 15) and requiring other outputs of mapping flag distribution (-g) and insert size histogram plot for each BAM library (-h). SVs were called using breakdancer-max on tumour and normal samples with the resulting configuration file and requiring allele frequency column in output file (-h). The output file (sv.out) was filtered for only somatic SVs with no supporting read (num_Reads) in matched normal sample.

*Commands used:*

[1] Create configuration file

```
bam2cfg.pl -q 15 -g -h ${TUMOR_BAM} ${NORMAL_BAM} > sv.cfg
```

[2] Call SVs with the configuration file from previous step

```
breakdancer-max -h sv.cfg > sv.out
```

#### CNVKit

CNVKit was used to call copy number in "wgs" mode, specifying the number of CPU cores (-p $CORE) and assuming male-reference (-y) as chromosome Y was included in SV simulation and CNV segments were converted to SVs in VCF format.

*Commands used:*

[1] Copy number calling pipeline

```
cnvkit.py batch $TUMOR_BAM --normal $NORMAL_BAM --method wgs \
--fasta ${REF_hg38} \
--annotate refFlat.txt -p ${CORE} \
--output-reference cnvkit_sv_ref.cnn -y –output-dir ${RUN_Dir}
```

### [2] Derive absolute integer copy number for each segment, using the output CNS file from previous step

```
cnvkit.py call ${TUMOUR_name}.cns -y -o ${TUMOUR_name}.call.cns
```

### [3] Convert to vcf file using output CNS and CNR files from previous steps

```
cnvkit.py export vcf ${TUMOUR_name}.call.cns –cnr ${TUMOUR_name}.cnr -i
"randomSV2" -o{vcf_name}.vcf
```

#### Pindel

First, a configuration file is created to specify the tumour and normal bam file locations and insert size of the sequencing library. Pindel was then run for each chromosome (e.g. chromosome 1; -c chr1) with 8 threads (-T 8), expected sequencing error rate (-e 0.02), maximum number of SVs to be detected (-x 1; 1= 128), bin size of reference (-w 1; 1=1 million) and requirement of at least four supporting reads (-M 4). Pindel generates an output file for each SV type (*_D for DEL, *_TD for DUP, *SI for INS, and *_INV for INV) for each chromosome. Each output file is then converted to VCF format with pindel2vcf, specifying name and version of the reference genome (- R ucsc.hg38), date of the version of the reference genome used (-d 122013), requirement of compatibility with GATK (-G) and at least four of supporting reads (-e 4). VCF files for each SV type per chromosome are then merged using vcfcat (https://github.com/vcflib/vcflib), resulting in one VCF file. Somatic SVs are filtered on the VCF file with no supporting reads (VCF field AD = 0) in matched normal sample.

*Commands used:*

[1] Call SVs in each chromosome (e.g. chromosome 1), generating an output file per SV type (e.g. pindel_chr1_D, pindel_chr1_TD, pindel_chr1_SI and pindel_chr1_INV).

```
pindel -f ${REF_hg38} -i $Configure_file -o pindel_chr1 \
-c chr1 -T 8 -e 0.02 -x 1 -w 1 -M 4
```

[2] Convert each output for each chromosome (e.g. DEL for chromosome 1; -p pindel_chr1_D) from previous step to VCF format

```
pindel2vcf -r ${REF_hg38} -p ${pindel_chr1_D} \
-R ucsc.hg38 -d 122013 -G -e 4
```

#### Lumpy

Lumpy was run following the recommended workflow (https://github.com/arq5x/lumpy-sv) by first generating discordant paired-end alignments and split-read alignments in the pre-processing step. Lumpy express was then executed jointly on tumour and normal samples with verbose mode (-v) for printing errors and warnings and at least one evidence (supporting reads) for a call (-m 1). The default minimum number of supporting reads is 4 set by Lumpy. Instead of using this threshold in SV calling, we use it in the following filtering step. Somatic SVs are filtered on the output VCF file with minimum of four supporting reads for each variant in the tumour sample (VCF field INFO/SU=4) and the variant allele not being observed in the matched-normal sample (VCF field INFO/SU=0).

*Commands used:*

[1] Pre-processing for both tumour and normal samples (example shown for one sample)

```
# Extract the discordant paired-end alignments
samtools view -b -F 1294 ${TUMOR_BAM} > tumour.discordants.bam
# Extract the split-read alignments
samtools view -h ${TUMOR_BAM} \
| lumpy-sv/scripts/extractSplitReads_BwaMem -i stdin \
| samtools view -Sb - \
> tumour.splitters.bam
# Sort both discordant paired-end and split-read alignments from
previous two steps
samtools sort -o tumour.discordants.sort.bam tumour.discordants.bam
samtools sort -o tumour.splitters.sort.bam tumour.splitters.bam
```

[2] Running Lumpy Express jointly on tumour and normal samples with inputs of pre-extracted sorted splitter and discordant alignments from pre-processing step and output of VCF file.

```
lumpyexpress -v -m 1 \
 -B ${TUMOR_BAM},${NORMAL_BAM} \
 -S tumour.splitters.sort.bam,normal.splitters.sort.bam \
 -D tumour.discordants.sort.bam,normal.discordants.sort.bam \
 -o ${vcf_name}.vcf
```

##### Delly

Delly SV calling was performed by *delly call*, with output SVs in BCF format, which were then filtered with *delly filter* in somatic filter mode (-f somatic) and "PASS" in the VCF FILTER field (-p). A tab-delimited sample description file (-s samples.tsv), where the first column is the sample id (as in the BCF file) and the second column is either 'tumor' or 'control', is required for somatic filtering by *delly filter*. The output BCF file from *delly filter* was then converted to VCF format.

*Commands used:*

[1] Call SVs and report in BCF format

```
delly call -o ${vcf_name}.bcf -g ${REF_hg38} ${TUMOR_BAM} ${NORMAL_BAM}
```

[2] Filter SVs from the BCF file from previous step

```
delly filter -p -f somatic -o ${vcf_name}.somatic.PASS.bcf -s
samples.tsv ${vcf_name}.bcf
```

##### Manta

Manta was run in two steps: somatic configuration to create the workflow run script and workflow execution in local run mode (-m local) and specifying number of cores to use (-j ${CORES}. SV calls were filtered with "PASS" in the VCF FILTER field in output VCF file.

*Commands used:*

[1] Configuration in tumour and normal analysis to create the workflow run script ${RUN_Dir}/runWorkflow.py

```
configManta.py --normalBam ${NORMAL_BAM} --tumorBam ${TUMOR_BAM} --
referenceFasta ${REF_hg38} --runDir ${RUN_Dir}
```

[2] Execution with the workflow run script from previous step

```
python ${RUN_Dir}/runWorkflow.py -m local -j ${CORES}
```

##### GRIDSS

GRIDSS variant calling pipeline was run with default options with tumour ($TUMOUR) and normal ($NORMAL) BAM files, specifying output VCF ($OUTPUT) and assembly ($ASSEMBLY) and a blacklist in BED format (BLACKLIST), shown in the following commands. ENDNOTE DAC blacklist is recommended by GRIDSS to exclude regions such as telomeric and centromeric regions from analysis. Output SVs were assigned SV types using simple-event-annotation.R script, which was included in the example folder of GRIDSS package. Somatic SVs were filtered based on VCF QUAL filed of zero for normal sample and further filtered with "PASS" in the VCF FILTER field for high-confidence SVs. Although high-confidence SVs can also be filtered based on VCF QUAL (e.g. QUAL ≥ 500), we used "PASS" criteria to be consistent with other SV callers and avoid defining the QUAL threshold.

*Command used:*

```
GRIDSS_JAR=gridss/gridss-1.8.1-gridss-jar-with-dependencies.jar
java -ea -Xmx31g \
        -Dsamjdk.create_index=true \
        -Dsamjdk.use_async_io_read_samtools=true \
        -Dsamjdk.use_async_io_write_samtools=true \
        -Dsamjdk.use_async_io_write_tribble=true \
        -Dsamjdk.compression_level=1 \
        -Dgridss.gridss.output_to_temp_file=true \
        -cp $GRIDSS_JAR gridss.CallVariants \
        TMP_DIR=tmp \
        WORKING_DIR=${RUN_Dir} \
        WORKER_THREADS=${CORES} \
        REFERENCE_SEQUENCE="$REFERENCE" \
        INPUT="$NORMAL" \
        INPUT="$TUMOUR" \
        OUTPUT="$OUTPUT" \
        ASSEMBLY="$ASSEMBLY" \
        BLACKLIST="$BLACKLIST" \
        2>&1 | tee -a gridss.$HOSTNAME.$$.log
```

##### SvABA

SvABA was run with tumour and matched normal samples, specifying the number of cores (-p $CORES) and the output VCF file ($OUTPUT.svaba.somatic.sv.vcf) was filtered by the caller on somatic status and PASS quality. SvABA only report breakend (BND) in output file, therefore the SV types were classified according to the orientation of BND and its mate BND reported in ALT field in the output VCF file. The ALT field of BND (ALT1) and its mate (ALT2) were analysed as follows for SV type classification. If ALT1 starts with "N[" and ALT2 ends with "]N", the SV type is assigned as DEL, where N is any nucleotide. If ALT1 ends with "]N" and ALT2 starts with "N[", the SV type is assigned as DUP/INS. If both ALT1 and ALT2 start with "N]" or end with "[N", the SV type is assigned as INV. If the BND and its mate are in different chromosomes, they are annotated as BND events.

*Command used:*

```
svaba run -t $TUMOR_BAM -n $NORMAL_BAM -p $CORES -a $OUTPUT -G $REF_hg38
```

### Evaluation of somatic SV calls

We require true positive (TP) SV call to meet two criteria: i) reported SV type must match the simulated type (**Table 2**), and ii) detected breakpoints must be within T bp from the simulated breakpoints (**Figure 8**). In the evaluation study by Kosugi et al., they required true positive SV call to have sufficient overlapping with true event [8]. Due to the limitation of this approach for certain SV types, which cannot be defined by genomic region such as INS, Kosugi et al. required TP INS having breakpoint within 200bp of true INS breakpoint, same as our criterion (ii). Another limitation to this approach is the requirement of the criteria to define sufficient overlapping [10]. Lee *et al*. used the same criterion (ii) as we have, but did not require SV calls to match type due to the ambiguity of SV type specification by different SV callers [10]. By contrast, Cameron et al. used the same criterion (ii) as we have and imposed an additional criterion requiring the SV size of detected SV different to that of true event by at most 25% [9]. Requirement for SV size similarity is not suitable for all SV types, such as BND for inter-chromosomal rearrangements, due to the lacking information of rearranged sequence length. Therefore, we only imposed the two TP criteria in this study.

To pass the first criterion, SV types were compared according to **Table 2** as explained in main text. To pass the second criterion, all breakpoint positions corresponding to an event must be within a prescribed breakpoint precision threshold (T). That is, all reported breakpoints must be no more than T bp from simulated breakpoint: $d1 < T$ and $d2 < T$ (**Figure 5**). In this study, we examined T values ranging from 0 to 2000 bp.

The total number of true positives is calculated as the number of calls in a callset satisfying both TP criteria. The number of false positives (FP) is the number of SV calls in a callset not satisfying either or both TP criteria. Following the confusion matrix, we define: sensitivity = TP / (TP + FN) and precision = TP / (TP + FP).

##### Evaluation of union and intersection callsets

Two SV calls were considered the same if they have matching SV type (**Table 2**) and their reported breakpoint positions are within a threshold of 200 bp of each other. The threshold was set to be the same as the breakpoint resolution threshold for TP criteria. Thus, the union callset of two SV callers is defined as SVs detected by either Caller_1 or Caller_2, while the intersection callset is defined as calls reported by both Caller_1 and Caller_2. In both cases, Caller_1 is the "dominant caller" such that any overlapping calls in the final callset is taken from the output from Caller_1 (including coordinates and SV types).

##### Evaluation of segmental duplication region

Segmental duplication (SegDup) regions were based on the genomicSuperDups [last updated 19/10/2014] table from UCSC Genome Browser. The SegDup is considered to collocate with an SV if (i) at least 1 bp of the SV interval overlap a SegDup for DEL, DUP and INV, or (ii) at least one BND of a DINS, FINS, or TRA is within a SegDup interval. By this definition, 201 of our simulated SVs are within SegDup regions. SVs called by SV callers whinin SegDup regions are similarly defined. Sensitivity and precision are separately calculated for SVs within and without SegDup regions.
