## Supplementary material for "Detection of somatic structural variants from short-read next-generation sequencing data": Table S1

**Table S1. F1 score across different SV types for every SV caller.**

| | DEL | DUP | INV | BND |
|---|---|---|---|---|
| **BreakDancer** | 0.415 | n/a | 0.00 | n/a |
| **CNVKit** | 0.007 | 0.064 | n/a | n/a |
| **Pindel** | 0.708 | 0.476 | 0.017 | n/a |
| **Lumpy** | 0.954 | 0.805 | 0.966 | 0.948 |
| **Delly** | 0.950 | 0.804 | 0.890 | 0.681 |
| **Manta** | 0.949 | 0.881 | 0.953 | 0.898 |
| **GRIDSS** | 0.911 | 0.833 | 0.945 | 0.899 |
| **SvABA** | 0.792 | n/a | 0.880 | 0.847 |

*n/a: structural variant type not reported by the SV caller
